# System-wide identification and prioritization of enzyme substrates by thermal analysis (SIESTA)

**DOI:** 10.1101/423418

**Authors:** Amir Ata Saei, Christian M. Beusch, Pierre Sabatier, Juan Astorga Wells, Alexey Chernobrovkin, Sergey Rodin, Katja Näreoja, Ann-Gerd Thorsell, Tobias Karlberg, Qing Cheng, Susanna L. Lundström, Massimiliano Gaetani, Ákos Végvári, Elias S.J. Arnér, Herwig Schüler, Roman A. Zubarev

**Author notes:** Correspondence and materials requests for materials should be addressed to A.A.S. and R.A.Z.

## Abstract

Despite the immense importance of enzyme-substrate reactions, there is a lack of generic and unbiased tools for identifying and prioritizing substrate proteins which are modulated in the structural and functional levels through modification. Here we describe a high-throughput unbiased proteomic method called System-wide Identification and prioritization of Enzyme Substrates by Thermal Analysis (SIESTA). The approach assumes that enzymatic post-translational modification of substrate proteins might change their thermal stability. SIESTA successfully identifies several known and novel substrate candidates for selenoprotein thioredoxin reductase 1, protein kinase B (AKT1) and poly-(ADP-ribose) polymerase-10 systems in up to a depth of 7179 proteins. Wider application of SIESTA can enhance our understanding of the role of enzymes in homeostasis and disease, open new opportunities in investigating the effect of PTMs on signal transduction, and facilitate drug discovery.

---

At least a third of all proteins possess enzymatic activity. One of the most comprehensive enzyme databases BRENDA comprises >9 million protein sequences and encompasses 6953 classes of enzyme-catalyzed reactions (http://genexplain.com/brenda/) ^1^. Many of these enzymes catalyze the modifications of protein substrates. Only in human genome, an estimated 1,089 non-metabolic enzymes are present ^2^, including for example more than 500 putative kinases. Transient modulation of protein post-translational modifications (PTMs) controls numerous cellular processes by inducing a host of downstream effects, such as changes in protein function, stability, interactions, hemostasis, localization and cellular diversification ^3^. Not surprisingly, mechanisms and kinetics of protein modifications have become a vibrant research area. However, there still has been a lack of methods for prioritization of the substrates undergoing structural changes in terms of the functional impact of the modifications ^4, 5^. Furthermore, an important aspect of PTM research is the characterization of enzyme-substrate associations, which is essential for our understanding of cell biology and disease mechanisms. Moreover, many high-throughput screening assays rely upon modified substrates as a readout. The lack of information on the physiological substrates of enzymes hampers the development of effective therapeutics, *e.g.* in Parkinson’s disease ^6^ and cancer ^7^.

Existing techniques used for identifying specific substrates are enzyme-specific, labor-intensive and often not straightforward. Such experiments include the use of genetic and pharmacologic perturbations ^8^, substrate-trapping mutants ^9^, affinity purification-mass spectrometry ^10^, utilizing peptide ^11^ or protein arrays ^12^, tagging the client proteins by substrate analogues using engineered enzymes ^13^ and peptide immunoprecipitation ^14^ or the use of sophisticated computational tools ^15^. Most of these techniques are specifically designed for a certain enzyme or enzyme class, which limits their applicability. Engineering enzymes can alter the biology of the system, potentially introducing a bias. Therefore, designing an unbiased, general, quick and proteome-wide method not involving artificial modification of the enzyme or substrate can prove to be a significant methodological advancement and a complement to above approaches.

Mass spectrometry based CEllular Thermal Shift Assay (MS-CETSA) or Thermal Proteome Profiling (TPP) is a recent method that can assess system-wide protein binding to small molecules, metabolites or nucleic acids by monitoring changes in protein thermal stability ^16, 17^. Since PTMs can also alter protein thermal stability, these methods can be potentially used to probe proteome-wide effects of PTMs. For example, Nordlund *et al.* have shown that phosphorylation leads to extensive intramolecular reorganization and stabilization of retinoblastoma-associated protein 1 (RB1) ^18^, while Savitski *et al.* have shown a correlation between phosphorylation and protein stability in mitosis ^19^. By employing CETSA with a Western blot readout at a single protein level, it has been shown that O-GlcNAcylation enhances stability of Nod2 protein ^20^. Huang et al. have recently developed a method called Hotspot Thermal Profiling that relates shifts in peptide melting temperature in response to site-specific phosphorylation sites (hotspots) ^5^. A very important assertion made in this work is that, the larger the shift, the more likely is the biological importance of a given PTM. Therefore, proteome-wide monitoring of the thermal stability changes in the cell lysate upon addition of a recombinant enzyme and a cosubstrate has the potential of not only revealing the enzyme substrates, but also prioritizing them according to the altered stability they demonstrate when undergoing modification. In this paper, substrate is the protein post-translationally modified by the enzyme, while cosubstrate refers to a molecule (such as NADPH, ATP and NAD) that participates in the enzymatic reaction.

In many cases, the concomitant protein-enzyme and protein-cosubstrate interactions can mask modification-specific thermal stability changes of the substrates. This problem is addressed in our method of System-wide Identification and prioritization of Enzyme Substrates by Thermal Analysis (SIESTA). SIESTA identifies *specific* thermal stability changes induced in substrate proteins by a combination of enzyme and cosubstrate as compared to the changes induced by either enzyme or cosubstrate alone (workflow in Fig. 1). The idea of specific response is borrowed from our methods of Functional Identification of Target by Expression Proteomics (FITExP) ^21^ and ProTargetMiner ^22^. In this approach, using orthogonal partial least squares-discriminant analysis (OPLS-DA) ^23^, protein T_m_ in “enzyme + cosubstrate” treatment can be contrasted with those in “control” (cell lysate incubated with vehicle), “enzyme”-treated lysate, and “cosubstrate”-treated lysate. Here, we apply SIESTA to three distinct enzymes, showing that this method can reveal known and putative substrates that change their stability upon modification in each system and rank them by the probability of having biological impact.

**Fig. 1.**
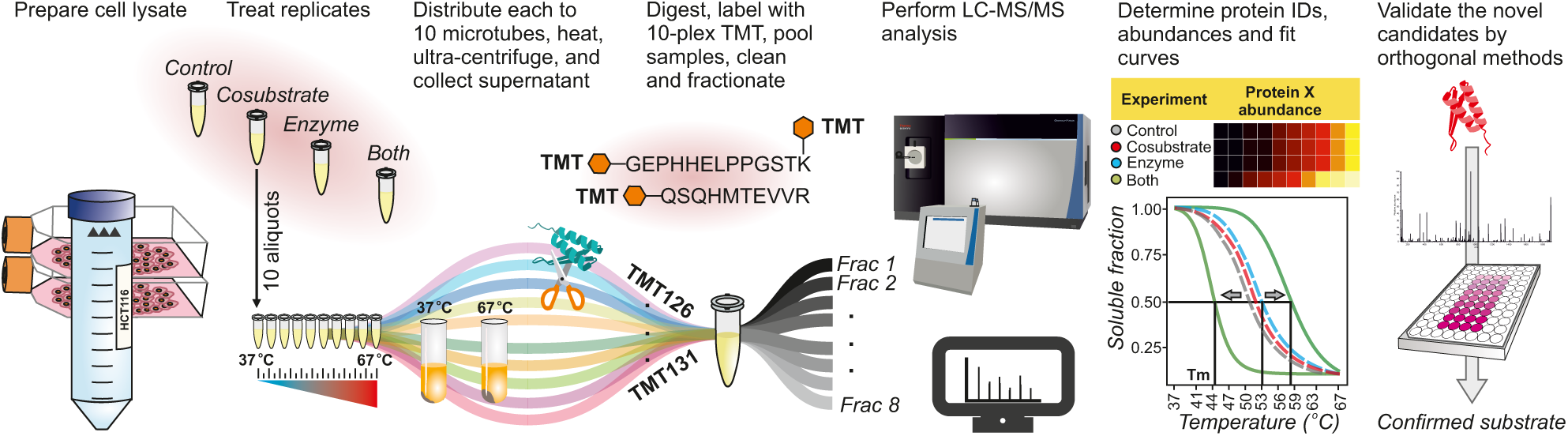
SIESTA workflow for unbiased proteome-wide identification of enzyme substrates. A master cell lysate is prepared by multiple freeze-thawing in a non-denaturing buffer. The cell lysate aliquots are treated with vehicle (control), cosubstrate, enzyme or combination of enzyme with cosubstrate (both). After treatment, each aliquot is split into 10 tubes, with each tube heated to a temperature point in the range from 37°C to 67°C. After removing unfolded proteins by ultracentrifugation, identical volumes of supernatants are digested with trypsin. The samples are then serially labeled with 10-plex TMT reagents, pooled, cleaned and fractionated by reversed-phase chromatography. After LC-MS/MS analyses of each fraction, protein IDs and abundances are determined, and sigmoid curves are fitted through an automated algorithm to determine the melting temperature T_m_ for each protein. For each non-vehicle treatment, the read-out is the protein’s ΔT_m_ shifts (of both signs) compared to control. Any protein shifting more upon addition of enzyme and cosubstrate compared to when they are added alone, are putative substrates of the enzyme under study. Such candidate protein substrates are subsequently confirmed by orthogonal verification methods.

## RESULTS

### SIESTA identified and ranked multiple known and putative TXNRD1 substrates

As the proof of principle, we selected an enzymatic reaction involving an oxidoreductase. Since such reduction reaction should destabilize substrate proteins and lead to negative ΔT_m_, the asymmetry between positive and negative values will be easy to verify. For this reaction we employed human selenoprotein thioredoxin reductase 1 (TXNRD1), a key oxidoreductase that catalyzes the reduction of specific substrate proteins using NADPH as a cosubstrate ^24^. A SIESTA experiment was performed in HCT116 cell lysate treated in duplicates with vehicle, NADPH, TXNRD1, or both (**Supplementary Data 1**).

Changes in T_m_ after NADPH treatment revealed stabilization of several known NADPH-interacting proteins (Fig. 2a), an example of which is shown in Fig. 2b. Among the 40 proteins annotated as NADPH binders in Uniprot database, 30 proteins (75%) were verified in our experiment, which indirectly validated the SIESTA approach (Fig. 2a). 247 novel proteins were identified as putative NADPH binders (**Supplementary Data 2**).

**Fig. 2.**
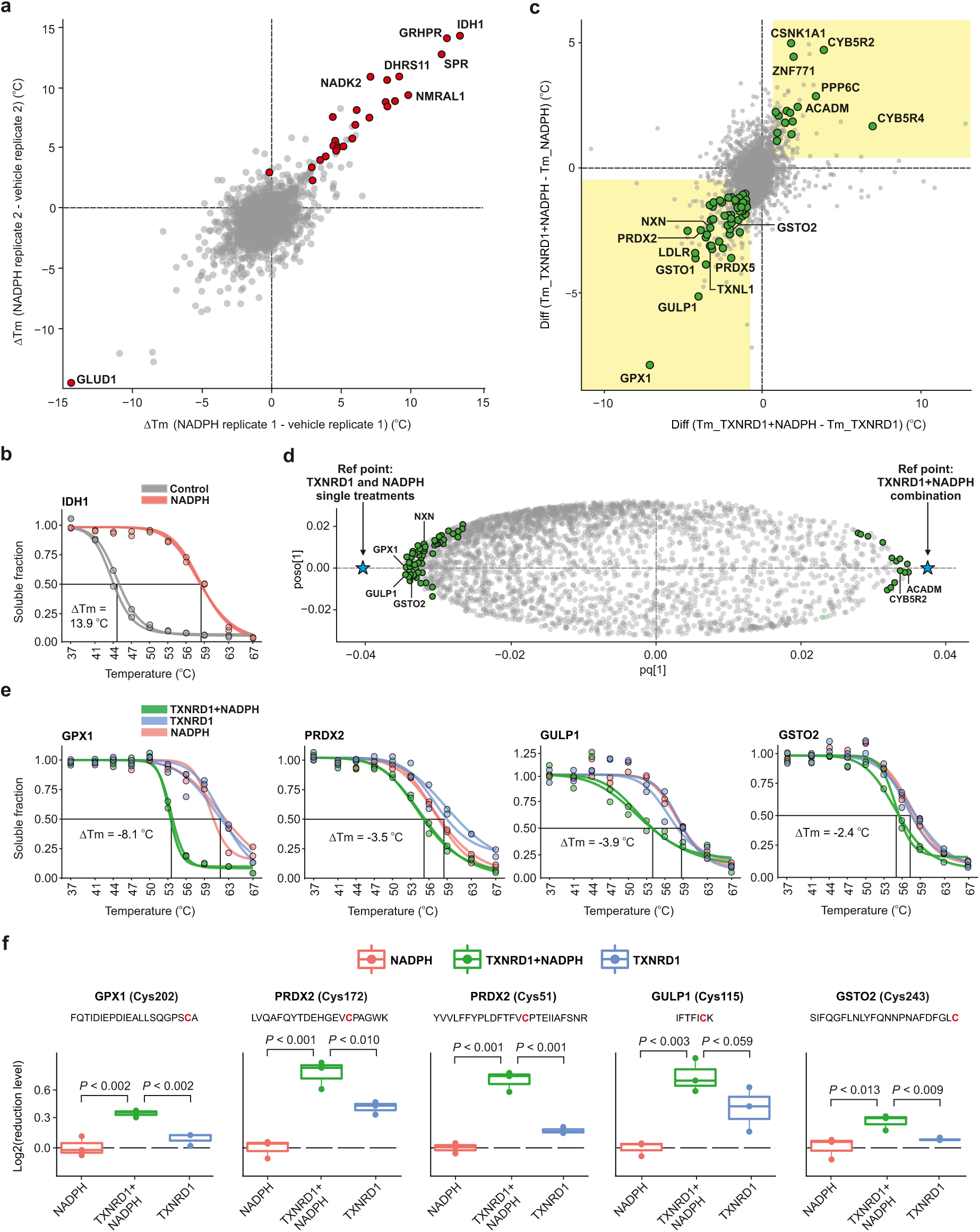
Proof-of-principle SIESTA experiment revealed known TXNRD1 substrates and suggested novel candidates. **a**, Scatterplot of protein T_m_ differences upon addition of NADPH in lysate. Known proteins from Uniprot are shown in red. **b**, Representative stabilization of NADPH binding protein IDH1. **c,** Scatterplot of T_m_ differences reveals the T_m_ shifts occurring only after simultaneous TXNRD1+NADPH addition; these shifts are thus likely due to enzymatic modifications (yellow shaded area, known and putative substrates are shown as green circles). **d**, Potential substrates (green circles) are mostly located close to the negative reference point (blue star) in an OPLS-DA model contrasting the “TXNRD1+NADPH” T_m_ against the single treatments. **e,** Representative melting curves of GPX1, PRDX2, GULP1 and GSTO2 are shown. **f**, Reduction of cysteines in the substrate proteins by incubation with TXNRD1+NADPH (*n*=3, one tailed Student t-test).

The analysis of specific ΔT_m_ shifts in the TXNRD1+NADPH treatment revealed that in the presence of NADPH, TXNRD1 destabilized both known and novel candidate substrate proteins (**Supplementary Data 3**). In general, the expected asymmetry in T_m_ shifts in favor of destabilization was well pronounced (Fig. 2c). An OPLS-DA model contrasting TXNRD1+NADPH with enzyme and cosubstrate single treatments was also used to reveal the specifically shifting proteins and rank them by their shifts and variable influence on projection (VIP)-values (Fig. 2d**, Supplementary Data 4**). In OPLS-DA, the VIP-values show the impact each x-variable (i.e. protein) have on the model with a higher value corresponding to a greater contribution ^22, 25^. In Supplementary Fig. 1a, the VIP-plot of the top 20 ranked proteins in the TXNRD1 OPLS-DA model are shown with the error bars representing 95% confidence intervals.

Examples of melting curves for proteins destabilized by TXNRD1 are shown in Fig. 2e. The 78 identified putative substrates (63 destabilized) mapped to the following INTERPRO Protein Domains and Features pathways: “Thioredoxin-like superfamily” (11 proteins, p < 1.3e-09) and “Thioredoxin domain” (7 proteins, p < 8.7e-08). The identification of new substrates for TXNRD1 is not surprising, as the mammalian TXNRD1 is known to have an easily accessible and highly reactive selenocysteine-containing active site ^26^.

GPX1 was the protein showing the strongest destabilization. Some GPX isoenzymes are known to be directly reduced by TXNRD1 ^27^. Among the identified TXNRD1 substrates, TXNL1 (or TRP32) ^28^ and NXN ^29^ are well known. Note that secondary reactions are unlikely during SIESTA, as the typical cellular volume is diluted ≈77 fold. It should also be noted that cell lysate is usually used for discovery of direct interactions in thermal profiling ^30^. Furthermore, if secondary reactions were possible, they would also occur in lysates treated with NADPH alone, and thus would be filtered away in our analysis.

To prove that the identified proteins can be directly reduced by TXNRD1, we designed a sequential iodoTMT labeling approach, with which the reduction/oxidation can be quantitatively analyzed on the single cysteine level. For this purpose we incubated the recombinant candidate proteins GPX1, GPX4, GSTO1, GSTO2, PRDX2, PRDX6 and GULP1 with TXNRD1+NADPH under the same conditions as in the SIESTA experiment. The results confirmed that GPX1, GPX4, GSTO2, PRDX2 and GULP1 can be directly reduced by TXNRD1 (Fig. 2f, Supplementary Fig. 1b for GPX4 and **Supplementary Data 5**). For example, in PRDX2 both Cys51 and Cys172, which form an interchain disulfide bond, were found reduced ^31^. GULP1 was reduced on Cys115 by TXNRD1. Interestingly, GULP1 exists as a dimer *in vivo* ^32^ and we noted the increased monomer levels for this protein upon incubation with TXNRD1+NADPH (Supplementary Fig. 1c-d). We however could not confirm the reduction of GSTO1 and PRDX6. Although this might be due to the absence of certain peptides in the MS data, one could estimate the false positive rate to be not higher than ∼30%. The fact that PRDX2 was detected here as a direct substrates for TXNRD1 showed that the enzyme has a capacity to also directly reduce these protein disulfides to some extent; alternatively traces of TXN present in the lysate could have been thought sufficient to facilitate this reaction because PRDX proteins are highly abundant. However, the validation redox proteomics experiment in Fig. 2f showed that TXNRD1+NADPH alone can indeed reduce PRDX2.

There were a number of proteins which were stabilized in the TXNRD1+NAPDH treatment, such as CYB5R2 and ACADM. This stabilization might be due to the protein interaction with the reduced form of TXNRD1, or by reduced species of these protein substrates forming other more thermostable states. TXN and TXNDC17, two known substrates of TXNRD1 ^33^, were absent in the SIESTA output due to their melting behavior. For example, although TXN was quantified in all replicates, it remained 63% soluble on average even at 67°C. Therefore, it was not possible to measure its T_m_ by fitting a sigmoid curve, and thus TXN was automatically excluded from analysis (TXN also did not melt well in PARP10 and AKT1 experiments in HCT116 and HELA cells, respectively). Thioredoxin reductase is also known to reduce GLRX2 and protein disulfide isomerase (PDI) in the literature. GLRX2 did not shift in our SIESTA experiment. We quantified all 6 PDIAs in our experiment, of which only PDIA6 was destabilized by −0.69°C and was therefore excluded by our criteria. Whether these two proteins are substrates of the human TXNRD1 is yet to be seen. Therefore, considering only TXN, TXNL1, NXN and TXNDC17 as known substrates of TXNRD1, SIESTA had a false negative rate of 50% in this system.

### SIESTA identified and ranked many novel putative substrates for protein kinase B (AKT1)

We decided to confirm the utility of SIESTA for phosphorylation as a ubiquitous and small modification. We chose the AKT1 (protein kinase B) as a model system due to its importance in metabolism, proliferation, cell survival, growth and angiogenesis. In AKT1 SIESTA experiment (data in **Supplementary Data 6**), ATP was used at 500 µM, at which concentration it only acts as a cosubstrate ^34^. 31% (123/396) of the proteins annotated in Uniprot as ATP binders were also verified in our experiment. 257 proteins were identified as novel putative ATP binders (Supplementary Fig. 2a, **Supplementary Data 7**). ACTB and MAP2K4 melting curves are shown as examples in Supplementary Fig. 2b. In total, 44 proteins were identified as putative AKT1 substrates (Fig. 3a, **Supplementary Data 8**), among which TRIP12, MEF2D, and BCL3 were known. Interestingly, BCL3 is known to be specifically modified and stabilized by AKT1 ^35^. The melting curves for representative substrates are shown in Fig. 3b. Among the remaining 41 proteins, 4 molecules were known to interact with AKT1 (CAMKK1, PLEKHF2, CEP76 and IMPDH2). In alternative methods, such as KISS ^36^, any protein interacting with a kinase is considered to be a potential substrate ^37^. OPLS-DA loading rankings of the putative substrates and their VIP-values are provided in **Supplementary Data 9 (**Supplementary Fig. 2c).

**Fig. 3.**
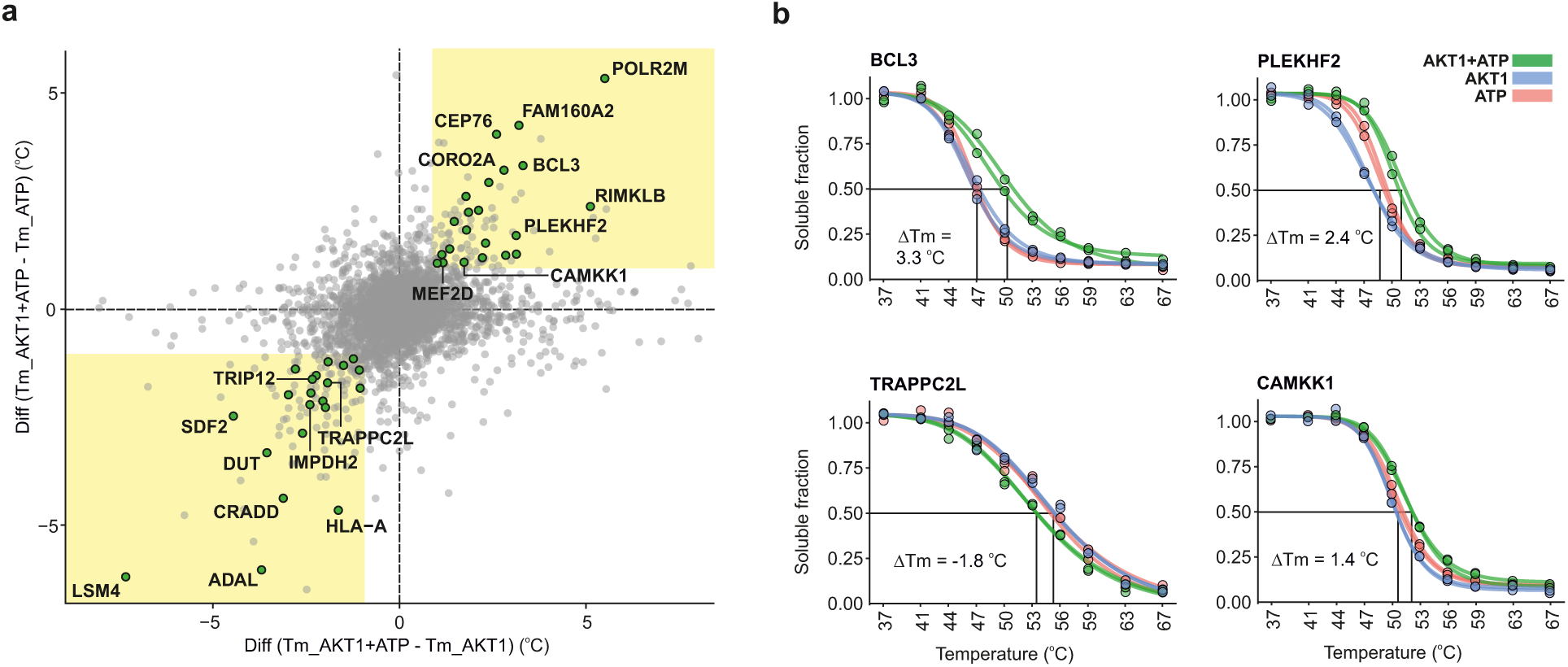
SIESTA identified known and putative substrates for AKT1 kinase. **a**, Scatterplot of T_m_ differences reveals the T_m_ shifts occurring only after simultaneous AKT1+ATP addition. **b,** Representative melting curves for known and putative AKT1 substrates.

To further validate a number of putative substrates, we incubated recombinant PLEKHF2 and TRAPPC2L with AKT1+ATP under the exact conditions as in SIESTA. PLEKHF2 is already known to interact with AKT1 ^38^. Tracking phosphate release from these recombinant proteins confirmed their modification by AKT1 (Supplementary Fig. 2d). We chose this approach over phosphoproteomics, as the former will detect phosphorylation present in *any* peptide belonging to the protein, and not only single phosphopeptides.

### SIESTA identified and ranked many novel putative substrates for PARP10

We next selected the poly-(ADP-ribose) polymerase-10 (PARP10) system that performs mono-ADP ribosylation of proteins ^39^. ADP-ribosylation is involved in cell signaling, DNA repair, gene regulation and apoptosis. Identification of PARP family substrates by mass spectrometry has generally proven challenging, as ADP-ribosylation is a glycosidic modification that can be easily lost during protein extraction or sample processing. It is also highly labile in the gas phase, which hampers its detection by MS/MS. Different strategies have thus been used to enrich the modified peptides for mass spectrometric analysis and use “gentle” MS/MS methods ^40, 41^. Although the identification of ADP-ribosylated substrates has been challenging for other techniques, since it is a large modification, it should be amenable to SIESTA.

As a verification of PARP10 SIESTA analysis (data in **Supplementary Data 10**), 22% (9/41) of proteins annotated as NAD binders were found, together with 87 putative new NAD binding proteins (Supplementary Fig. 3a, listed in **Supplementary Data 11**). CTBP2 and GALE are shown as known examples in Supplementary Fig. 3b. In total, 58 proteins were identified as potential PARP10 substrates (Fig. 4a, **Supplementary Data 12**), some of which already known, such as ILF2, ILF3, IPO4 and PUM1 ^42^ as well as GAPDH ^43^. Melting curves for some of the putative substrates are shown in Fig. 4b. An OPLS-DA model contrasting “PARP10+NAD” T_m_ *vs.* those from all other treatments is given in Supplementary Fig. 3c, and the OPLS-DA loading rankings of the putative substrates and their VIP values can be found in **Supplementary Data 13**.

**Figure 4.**
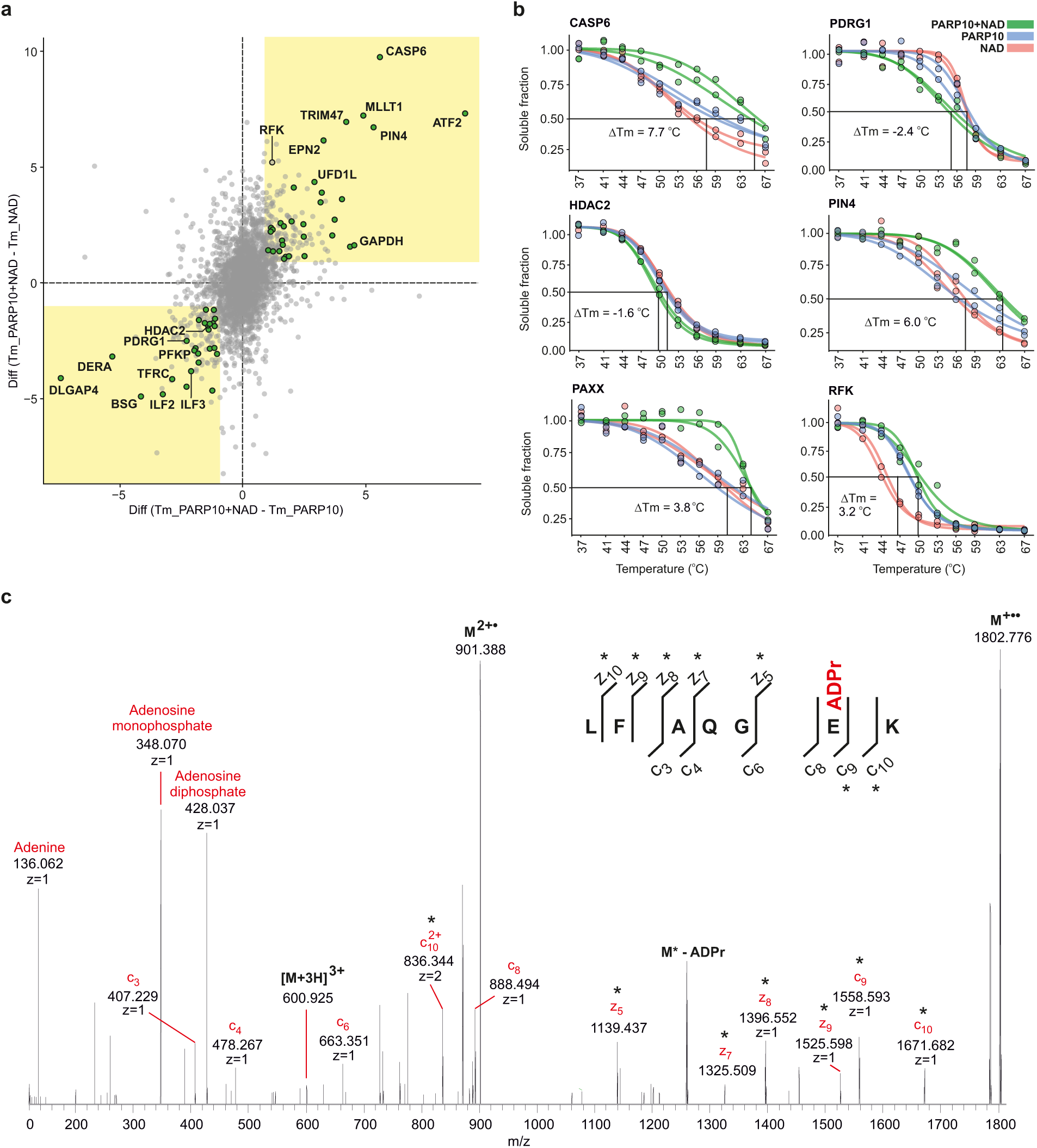
SIESTA identified and ranked known and novel PARP10 substrates. **a**, Scatterplot of T_m_ differences reveals the shifts (shown with green circles) occurring when PARP10+NAD are incubated with cell lysate. **b,** Representative melting curves of putative PARP10 substrates. **c,** Mono-ADP-ribosylation on a glutamic acid residue Glu110 in the PDRG1 peptide with the highest sequence-fitting score revealed by targeted ETD MS/MS of 3+ molecular ions. The fragments carrying the modification are marked with an asterisk.

The majority of the identified PARP10 substrates were novel, reflecting the limited number of previous studies in this area. We used targeted mass spectrometry to verify the PARP10-mediated mono-ADP-ribosylation of destabilized PDRG1 and HDAC2 as well as the stabilized PIN4 and CASP6, based on OPLS-DA rankings and availability of full-length recombinant proteins. We also validated RFK, as it showed a dramatic stabilization when compared to NAD and weak stabilization when compared to the PARP10 treatments. After incubation with recombinant PARP10 and NAD, the above proteins were digested and analyzed with LC-MS/MS. Every higher energy collision dissociation (HCD) MS/MS event triggered in data-dependent acquisition was investigated in real time for the presence of signature ions of adenine (*m/z* 136.0623), adenosine-18 (*m/z* 250.094) and adenosine monophosphate (AMP, *m/z* 348.0709). The presence of these triggers would then initiate a second MS/MS event using electron-transfer dissociation (ETD) with a supplementary HCD activation. The obtained PDRG1 sequence coverage was 74%, and the protein was found modified with ADP in three locations: on Glu110, Glu75 and Asp32 (Supplementary Table 1). The ETD MS/MS spectrum of a peptide with Glu110 is shown in Fig. 4c and the other sites are shown in Supplementary Fig. 4a-b. The RFK sequence coverage was 94%, and ADP ribose moieties were found in three positions: on Glu140, Glu131 and Glu113, ordered from highest to lowest peptide score (Supplementary Table 1). The ETD MS/MS spectrum of the peptide with highest score is shown in Supplementary Fig. 3d and the other sites are shown in Supplementary Fig. 4c-d. For HDAC2 and PIN4, the sequence coverage with trypsin digestion was not complete. Consequently, the ADP-ribosylation of HDAC2, PIN4 as well as PDRG1 and RFK was verified using a chemiluminescence assay (Supplementary Fig. 3e).

CASP6 (caspase-6) showed the strongest specific stabilization (ΔTm of 7.7°C, Fig. 4a,b), but its modification was not verified in either of the two *in vitro* assays. It should also be noted that PARP10 was suggested to be a substrate for caspase-6 during apoptosis ^44^. PARP10 has a major cleavage site at Asp406 that is preferentially recognized by caspase-6 ^44^. The strong specific thermal stabilization might therefore indicate that PARP10 induces a conformational change in caspase-6 and thus an increase in its stability by binding, as has been reported for other caspase-6 substrates ^45^. The reason why caspase-6 stabilization was not observed upon PARP10 addition in the absence of NAD is that auto-modified PARP10 is required for effective caspase-6 binding ^44^. With four out of five proteins being validated as PARP10 substrates (except for caspase-6), the false positive rate of SIESTA would be 20% in this system.

## DISCUSSION

We demonstrate SIESTA to be a general approach for unbiased identification and prioritization of functional protein substrates for specific enzymes in a proteome-wide manner. We uncovered a number of known or novel candidates for TXNRD1, AKT1 and PARP10 enzymes, implicating them in important cellular processes. Applying the “HotSpot” ideology to rank the putative substrates, SIESTA can be useful in uncovering the biophysical consequences of PTMs, as it can provide the most plausible substrates for functional validation. Besides the use in fundamental research, SIESTA can also facilitate drug development by discovery of substrates that can be used in screening for enzyme inhibitors.

Here we show the applicability of SIESTA for three distinct enzymes, and the utility of SIESTA for other enzyme systems will have to be established in further studies. It can probably be hypothesized that any modification will have some effect on protein stability, even though some of these changes may be too minute to be detected. Such substrates can be likely discovered via improving the statistical power, e.g., by adding more biological replicates.

While SIESTA will discover only substrates that significantly change their thermal stability upon modification, according to HotSpot conjecture such substrates are more likely to be biologically relevant. This makes it difficult to compare SIESTA results with those obtained by other methods that typically lack such ranking ability. Technically, the SIESTA, false positive substrate discovery rate for TXNRD1 and PARP10 enzymes was around 30 and 20%, respectively. Since different PTMs induce thermal shifts of different magnitudes, the false-positive and -negative rates will be dependent on the enzyme under study. The false negative rate for TXNRD1, the enzyme inducing large shifts, could be estimated as ≈50% (this is while we identified 78 substrates, compared to 4 known substrates in the literature), while the value must be higher for the AKT1 system, as majority of phosphorylation events do not have large impact on protein stability ^46^. It must however be noted that we used a 1°C ΔT_m_ minimum cutoff for selection of substrates. Setting a smaller cutoff, especially for phosphorylation as a small modification, will increase the number of identified substrates. For example, reducing the cutoff to 0.5°C increases the number of AKT1 substrates by 28, adding 5 more known substrates (TBC1D4, PLK1, PDE3A, SSBP4 and AKT1 itself). It should also be considered that phosphorylation is a dynamic modification, and the experimental conditions and choice of cell lines affects the list of substrates. Furthermore, not all proteins or phosphopeptides are detected in all experiments. This is why the lists of AKT1 substrates revealed in different studies have only few overlaps. For example, papers that have performed direct screening of AKT1 substrates in human cells ^47, 48^, only have 6 overlapping substrates with PhosphoSitePlus database ^49^.

Unlike the original HotSpot approach that compares the shifts of individual modified peptides with those of the whole bulk protein, thus identifying modifications that may or may not have significant occupancies, SIESTA compares the shift of the whole bulk protein with and without the enzyme and the cofactor, thus requiring the majority of protein molecules to be modified to be identified as substrate. Therefore, highly ranked SIESTA hits are less likely to fail in subsequent functional validation.

The spatial resolution of the method can be increased by sub-cellular fractionation of the lysate prior to analysis. Furthermore, cell-or tissue-specific substrates should be possible to discover by comparing lysates from different sources. Since the addition of enzyme in excess can cause non-physiological modifications, the identified candidates should be validated by other techniques, as suggested in Fig. 1. Mild detergents such as NP40 can be used to increase the representation of membrane proteins in SIESTA ^50^. On the other hand, the number of missing values can be reduced by using our high-throughput approach to thermal profiling, Proteome Integral Solubility Alteration (PISA) assay ^51^. Affinity purification approaches have the advantage of enriching for low abundant and low stoichiometry substrates. However, such methods suffer from missing the weak binders. This gap can be filled by SIESTA.

The studies focused on investigating the effect of PTMs on protein stability are scarce ^52, 53^ and mostly performed on a single protein level. How minuscule changes in chemical structure can lead to conformational and stability changes and manifest signaling and phenotypic consequences, is not fully explored. For example, it has been shown that glycosylation sites on CUB1 domain of Bone Morphogenetic Protein-1 (BMP1) are important for thermal stability and secretion of this protein ^52^. Even small modifications such as deamidation can change protein stability against temperature and urea ^54^. Therefore, one of the main application of SIESTA will be to study such events and decipher the biological roles of such modifications in a proteome-wide manner.

Generalization of the effect of PTMs on protein folding and stability can be difficult, since specific protein-PTM contacts do not necessarily follow general rules and might have evolved to confer beneficial energetic effects on protein folding ^53^. For example, N-glycosylation is known to generally increase the stability of target substrates ^55^, but random introduction of N-glycans in a protein does not necessarily stabilize the protein significantly. The interpretation of disulfide bond reduction on protein stability is more straightforward. However, the impact of other PTMs such as ADP-ribosylation and phosphorylation on protein stability might be harder to understand. Conformational distortion by addition of modification ^56^, conformational entropy and free energy ^57^ and size and position of modification ^58^ as well as changes in the charge state and solvent accessibility determine the outcome of a modification on overall protein stability ^58^. Furthermore, multiple modifications might have cumulative effects on the stability of a protein ^59^. It has also been suggested that the final stability of a given protein is governed the detailed energetics of the proteoforms ^60^.

Summarizing, the ease, breadth and speed of identifying enzyme-specific substrates offered by SIESTA can enhance our understanding of enzyme systems and disease, accelerate constructing high-throughput assays and thus facilitate drug discovery.

## METHODS

### Cell culture

Human colorectal carcinoma HCT116 (ATCC, USA) cells were grown at 37°C in 5% CO_2_ using McCoy’s 5A modified medium (Sigma-Aldrich, USA) supplemented with 10% FBS superior (Biochrom, Berlin, Germany), 2 mM L-glutamine (Lonza, Wakersville, MD, USA) and 100 units/mL penicillin/streptomycin (Gibco, Invitrogen). Human A549 cells (ATCC, USA) were grown under the exact same conditions in DMEM. Low-number passages (<10) were used for the experiments.

### Recombinant proteins

Human TXNRD1, GPX1 and GPX4 were expressed recombinantly in *E. coli* and purified as described earlier ^61^. PARP10 full length protein (used in SIESTA) and catalytic domain construct (used in validation assays) were produced as detailed before ^62^. The rest of the recombinant proteins were purchased and are listed in Supplementary Table 2.

### SIESTA experiment

Cells were cultured in 175 cm^2^ flasks, and were then trypsinized, washed twice with PBS, resuspended in 50 mM HEPES pH 7.5, 2 mM EDTA (for TXNRD1) or in 50 mM HEPES pH 7.5, 100 mM NaCl and 4 mM MgCl_2_ (for PARP10), both with complete protease inhibitor cocktail (Roche). For AKT1 experiment, phosphatase inhibitors (PhosSTOP, Sigma) were also added. Cells were lysed by five freeze-thaw cycles. The cell lysates were centrifuged at 21,000 *g* for 20 min and the soluble fraction was collected. The protein concentration in the lysate was measured using Pierce BCA assay (Thermo) and equally distributed into 8 aliquots (1 mL each). For TXNRD1, each pair of samples were incubated with vehicle, 1 mM NADPH, 1 µM TXNRD1, or with TXNRD1+NADPH at 37°C for 30 min. For PARP10, each pair of samples were incubated with vehicle, 100 µM NAD, 400 nM PARP10, or with PARP10+NAD at 37°C for 1 h. For AKT1, each pair of samples were incubated with vehicle (AKT1 buffer), 500 µM ATP (CAT# GE27-2056-01, sigma), 500 nM AKT1 or with AKT1+ATP at 37°C for 30 min. Each replicate was then aliquoted into 10 PCR microtubes and incubated for 3 min in SimpliAmp Thermal Cycler (Thermo) at temperature points of 37, 41, 44, 47, 50, 53, 56, 59, 63, and 67°C. Samples were cooled for 3 min at room temperature and afterwards kept on ice. Samples were then transferred into polycarbonate thickwall tubes and centrifuged at 100,000 g and 4°C for 20 min.

The soluble protein fraction was transferred to new Eppendorf tubes. Protein concentration was measured in the samples treated at lowest temperature points (37 and 41°C) using Pierce BCA Protein Assay Kit (Thermo), the same volume corresponding to 50 µg of protein at lowest temperature points was transferred from each sample to new tubes and urea was added to a final concentration of 4M. Dithiothreitol (DTT) was added to a final concentration of 10 mM and samples were incubated for 1 h at room temperature. Subsequently, iodoacetamide (IAA) was added to a final concentration of 50 mM and samples were incubated in room temperature for 1 h in the dark. The reaction was quenched by adding an additional 10 mM of DTT. Proteins were precipitated using methanol/chloroform. The dry protein pellet was dissolved in 8M urea, 20 mM EPPS (pH=8.5) and diluted to 4M urea. LysC was added at a 1 : 100 w/w ratio at room temperature overnight. Samples were diluted with 20mM EPPS to the final urea concentration of 1M, and trypsin was added at a 1 : 100 w/w ratio, followed by incubation for 6 h at room temperature. Acetonitrile (ACN) was added to a final concentration of 20% and TMT reagents were added 4x by weight to each sample, followed by incubation for 2 h at room temperature. The reaction was quenched by addition of 0.5% hydroxylamine. Samples were combined, acidified by TFA, cleaned using Sep-PaK cartridges (Waters) and dried using DNA 120 SpeedVac™ Concentrator (Thermo). The SIESTA samples for TXNRD1 and PARP10 were then resuspended in 0.1% TFA and fractionated into 8 fractions using Pierce™ High pH Reversed-Phase Peptide Fractionation Kit (Thermo). The AKT1 samples were resuspended in 20 mM ammonium hydroxide and separated into 96 fractions on an XBrigde BEH C18 2.1×150 mm column (Waters; Cat#186003023), using a Dionex Ultimate 3000 2DLC system (Thermo Scientific) over a 48 min gradient of 1-63% B (B=20 mM ammonium hydroxide in acetonitrile) in three steps (1-23.5% B in 42 min, 23.5-54% B in 4 min and then 54-63%B in 2 min) at 200 µL/min flow. Fractions were then concatenated into 24 samples in sequential order (*e.g.* 1,25,49,73).

### Sequential iodoTMT labeling

The proteins (2 µg each, in triplicates) were incubated with 1 mM NADPH, 1 µM TXNRD1, or with TXNRD1+NADPH at 37°C for 30 min. After solubilization in methanol, 4.4 mmol/L of iodoTMT was added to the samples (labels 126, 127 and 128 to replicate 1, 2 and 3 in each treatment) and incubated for 1h at 37°C with vortexing in the dark (free SH and SSH groups will be blocked in this stage). The proteins were precipitated using methanol chloroform and after drying, samples were dissolved in Tris buffer with 1% SDS and incubated at 37°C in the dark with 1 mM DTT for 1h. Subsequently, the samples were incubated with 4.4 mmol/L of the second iodoTMT label at 37°C in the dark for 1h (labels 129, 130 and 131 to replicates 1, 2 and 3 in each treatment). The reaction was quenched by 20 mM final concentration of DTT. NADPH, TXNRD1 and TXNRD1+NADPH-treated samples were then individually pooled and precipitated. Protein pellets were dissolved in Tris and urea 8M. The samples were then diluted to 4M urea, and lysC was added at a ratio of 1:100 enzyme: protein overnight. After dilution of urea to 1M, trypsin was added at a ratio of 1:100, followed by incubation for 6 h at 37°C. Samples were acidified by TFA and cleaned using SepPak and lyophilized using a vacuum concentrator. Samples were dissolved in 0.1% FA and 1 µg of each samples was analyzed with a Q Exactive instrument using a 2 h gradient.

### LC-MS/MS

After drying, samples were dissolved in buffer A (0.1% formic acid and 2% ACN in water). The TXNRD1 and PARP10 samples were loaded onto a 50 cm EASY-Spray column (75 µm internal diameter, packed with PepMap C18, 2 µm beads, 100 Å pore size) connected to the EASY-nLC 1000 (Thermo) and eluted with a buffer B (98% ACN, 0.1% FA, 2% H_2_O) gradient from 5% to 38% of at a flow rate of 250 nL/min. The eluent was ionized by electrospray, with molecular ions entering an Orbitrap Fusion mass spectrometer (Thermo).

The AKT1 samples were loaded with buffer A onto a 50 cm EASY-Spray column connected to a nanoflow Dionex UltiMate 3000 UHPLC system (Thermo) and eluted in an organic solvent gradient increasing from 4% to 26% (B: 98% ACN, 0.1% FA, 2% H_2_O) at a flow rate of 300 nL/min over 95 min.

The iodoTMT labeled samples were loaded with buffer A onto a 50 cm EASY-Spray column connected to an EASY-nLC 1000 (Thermo) and eluted with a buffer B (98% ACN, 0.1% FA, 2% H_2_O) gradient from 4% to 35% of at a flow rate of 300 nL/min over 120 min. The MS parameters of all the above mentioned experiments as well as the number of quantified proteins are summarized in Supplementary Table 3. An analysis of the total number of proteins and the number of proteins with missing values in different replicates in each SIESTA experiment are shown in Supplementary Fig. 5.

### Data processing

The raw LC-MS data (SIESTA) were analyzed by MaxQuant, version 1.5.6.5 ^63^. The Andromeda search engine matched MS/MS data against the Uniprot complete proteome database (human, version UP000005640_9606, 92957 entries). TMT10-plex on the MS/MS level was used for quantification of protein abundances. Cysteine carbamidomethylation was used as a fixed modification, while methionine oxidation was selected as a variable modification. In the AKT1 experiments, phosphorylation on serine and threonine was selected as variable modification, and used in quantification. For sequential iodoTMT labeling, TMT6-plex on the MS/MS level was used for quantification of peptide/protein abundances. Methionine oxidation was selected as a variable modification and a customized .fasta file with recombinant protein sequences was used. Trypsin/P was selected as enzyme specificity. No more than two missed cleavages were allowed. A 1% false discovery rate was used as a filter at both protein and peptide levels. For all other parameters, the default settings were used. After removing all the contaminants, only proteins with at least two peptides were included in the final dataset.

### Network mapping

For pathway analyses, STRING version 10.5 (http://string-db.org) protein network analysis tool was used with default parameters ^64^.

### Validation of mono-ADP-ribosylation by targeted tandem mass spectrometry

Recombinant RFK (5 µg) and PDRG1 (5 µg) were diluted with 50 Mm HEPES (pH = 7.5), 0.5 mM TCEP, 100 mM NaCl, 100 µM NAD, 4 mM MgCl_2_ and incubated with 400 nM of PARP10 for 1 h. Proteins were reduced with 10 mM DTT for 30 min and alkylated with 50 mM IAA for 30 min in the dark. Afterwards, 1M urea was added to the samples and LysC (overnight) and Trypsin (6 h) were added sequentially at 1 : 100 w/w to protein. After acidification, samples were cleaned using StageTips. Samples were dissolved in 0.1% FA and 1 µg of each samples was analyzed with LC-MS using a 1 h gradient.

The chromatographic separation of peptides was achieved using a 50 cm Easy C18 column connected to an Easy1000 LC system (Thermo Fisher Scientific). The peptides were loaded onto the column at a flow rate of 1000 nL/min, and then eluted at 300 nL/min for 50 min with a linear gradient from 4% to 26% ACN/0.1% formic acid. The eluted peptides were ionized with electrospray ionization and analyzed on an Orbitrap Fusion mass spectrometer. The survey mass spectrum was acquired at the resolution of 120,000 in the m/z range of 300-1750. The first MS/MS event data were obtained with a HCD at 32% excitation for ions isolated in the quadrupole with a *m/z* width of 1.6 at a resolution of 30,000. Mass trigger filters targeting adenine, adenosine and AMP ions were used to initiate a second MS/MS event using ETD MS/MS with HCD supplementary activation at 30% collision energy and with a 30,000 resolution. Samples treated with NAD but no PARP10 were used as negative controls.

Spectra were converted to Mascot generic format (MGF) using in-house written RAWtoMGF v. 2.1.3. The MGFs files were then searched against the UniProtKB human database (v. 201806), which included 71,434 sequences. Mascot 2.5.1 (Matrix Science) was used for peptide sequence identification. Enzyme specificity was set to trypsin, allowing up to two missed cleavages. C, D, E, K, N, R and S residues were set as variable ADP-ribose acceptor sites. Carbamidomethylation was set as a fixed modification on C and oxidation as a variable modification on M.

### *In vitro* mono-ADP-ribosylation assay

Hexahistidine-tagged PARP10 catalytic domain (auto-modification) or protein substrate (substrate protein modification) was immobilized on Ni^2+^-chelating microplates (5-PRIME). TEV-cleaved PARP10 catalytic domain was used for evaluation of substrate protein modification. Mono-ADP-ribosylation was assessed after incubation with 100 µM NAD^+^ (including 2% biotinylated NAD^+^, Trevigen) prior to chemiluminescence detection of biotinyl-ADP-ribose in a Clariostar microplate reader (BMG Labtech) as described in detail before ^62^.

### Phosphoprotein Phosphate Estimation Assay

The recombinant proteins were incubated with AKT1 and ATP under exact condition of SIESTA experiment. Samples treated with only AKT1 or ATP were used as controls. Phosphate release from the proteins was measured by the Phosphoprotein Phosphate Estimation Assay Kit (Thermo) according the manufacturer instructions. The absorbance was measured at 620 nm using an Epoch Microplate Spectrophotometer (BioTek).

### SDS-PAGE

GULP1 (3 µg) was incubated with NADPH (1 mM), TXNRD1 (1 µM) or their combination for 30 min at 37°C in triplicates. After addition of the NuPAGE LDS Sample Buffer (Thermo), the samples were loaded in a NuPAGE Bis-Tris Mini Gel (Thermo) with 10 lanes and separated on a NuPAGE Bis-Tris 4–12% gel in MOPS SDS Running Buffer under non-reducing conditions at 200V for 60 min using the XCell SureLock system (Thermo). SeeBlue Plus2 Pre-stained Protein Standard (Thermo) was used as a ladder. The gel was then washed and stained with Coomassie blue for 1 h and then destained overnight. The resulting protein bands were captured using Universal Hood II (Bio-Rad) and analyzed using Quantity One 4.6.9.

### Statistical Analysis

Curve normalization and fitting was done by an in-house R package (https://github.com/RZlab/SIESTA). Briefly, after removing the contaminant proteins and those quantified with less than two peptides, protein abundances in temperature points 41-67°C were normalized to the total proteome melting curve similar to Franken at al. ^30^. For each protein in each replicate, a sigmoid curve was fitted using non-linear least squares method according the formula:

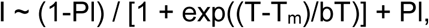

where Pl – high-temperature plateau of the melting curve, T_m_ – melting temperature, b – slope of the curve.

P values for the potential cosubstrate binding proteins were calculated by t-test based on the T_m_ values between vehicle and cosubstrate replicates. For selection of putative cosubstrate-binding proteins, the following criteria were used: 1) R2 > 0.7 between the measurement and the fitted curve, 2) the standard deviation between the replicates was <2.5°C, 3) P value < 0.05, 4) the absolute mean ΔTm larger than 1°C between the cosubstrate-and vehicle-treated samples.

P values for the potential substrates were calculated by two methods. In the first approach, t-test was made comparing the protein T_m_ values in Enzyme-Cosubstrate treatment against Enzyme and Cosubstrate single treatments. For selecting the putative substrates, the following criteria were used: 1) R2 > 0.7 between the measurement and the fitted curve, 2) the standard deviation between the replicates was <2.5°C, 3) p values between the Enzyme-Cosubstrate treatment against Enzyme and Cosubstrate treatments < 0.05 for one condition and <0.1 for the other; 4) the absolute mean ΔT_m_ was larger than 1°C for both conditions (a similar approach was used for selection of cosubstrate binding proteins).

Multivariate modeling using OPLS-DA was performed using SIMCA 15.0. Protein loading scores were validated using the VIP values at 95% confidence.

Two-tailed Student t-test (with equal or unequal variance depending on F-test) was applied to calculate p values, unless otherwise specified.

### Code availability

The curve fitting R package is available in GitHub (https://github.com/RZlab/SIESTA) with no access restrictions.

### Data availability

Excel files containing the analyzed data are provided in Supplementary Materials. The mass spectrometry data were deposited to the ProteomeXchange Consortium (http://proteomecentral.proteomexchange.org) via the PRIDE partner repository ^65^ with the dataset identifiers PXD010554 (for PARP10 and TXNRD1) and PXD014445 (for AKT1).

## ACKNOWLEDGEMENTS

We are grateful to Marie Ståhlberg and Carina Palmberg for their assistance in different proteomic experiments. The research is funded by grants from the Knut and Alice Wallenberg Foundation (grant KAW 2015.0063) and VINNOVA (grant Oxidocurin) awarded to R.Z.; E.A. is supported by grants from Karolinska Institutet, Swedish Cancer Society and Swedish Research Council, and H.S. by Swedish Research Council (grant 2015-4603). K.N. was supported by a stipend from the Wenner-Gren foundation. A.A.S. was supported by the gold level award from the Thermo Scientific Tandem Mass Tag Research Awards.

## AUTHOR CONTRIBUTIONS

Conceptualization, R.A.Z. and A.A.S.; methodology and experiment design, A.A.S., R.A.Z., C.M.B, J.A.W., H.S., E.A., S.R. and T.K.; project organization, training, resources and funding acquisition, R.A.Z., H.S. and E.A; SIESTA experiments, A.A.S., P.S. and M.G.; redox experiments, A.A.S. and P.S.; targeted mass spectrometry for ADP-ribose confirmation, A.A.S. and A.V.; data analysis and visualization, A.A.S., C.M.B., S.L.L. and A.C.; production and testing of enzymes, A.G.T. and Q.C.; *in vitro* mono-ADP-ribosylation assays, K.N. and H.S.; writing—original draft, A.A.S. and R.A.Z.; writing—review & editing, A.A.S., R.A.Z., H.S., E.A and J.A.W.

## Supplementary Data (Excel tables)

**Supplementary Data 1.** The processed data for TXNRD1 SIESTA experiment for all the replicates

**Supplementary Data 2.** NADPH binding proteins

**Supplementary Data 3.** The identified substrates for TXNRD1

**Supplementary Data 4.** OPLS-DA scores contrasting TXNRD1+NADPH vs. TXNRD1 and NADPH single treatments

**Supplementary Data 5.** The reduced ratio of peptides from candidate substrate proteins in the presence of TXNRD1, NADPH or TXNRD1+NADPH

**Supplementary Data 6.** The processed data for AKT1 SIESTA experiment for all the replicates

**Supplementary Data 7.** ATP binding proteins

**Supplementary Data 8.** The identified substrates for AKT1

**Supplementary Data 9.** OPLS-DA scores contrasting AKT1+ATP vs. AKT1 and ATP single treatments

**Supplementary Data 10.** The processed data for PARP10 SIESTA experiment for all the replicates

**Supplementary Data 11.** NAD binding proteins

**Supplementary Data 12.** The identified substrates for PARP10

**Supplementary Data 13.** OPLS-DA scores contrasting PARP10+NAD vs. PARP10 and NAD single treatments

## Supplementary information

**Supplementary Fig. 1.**
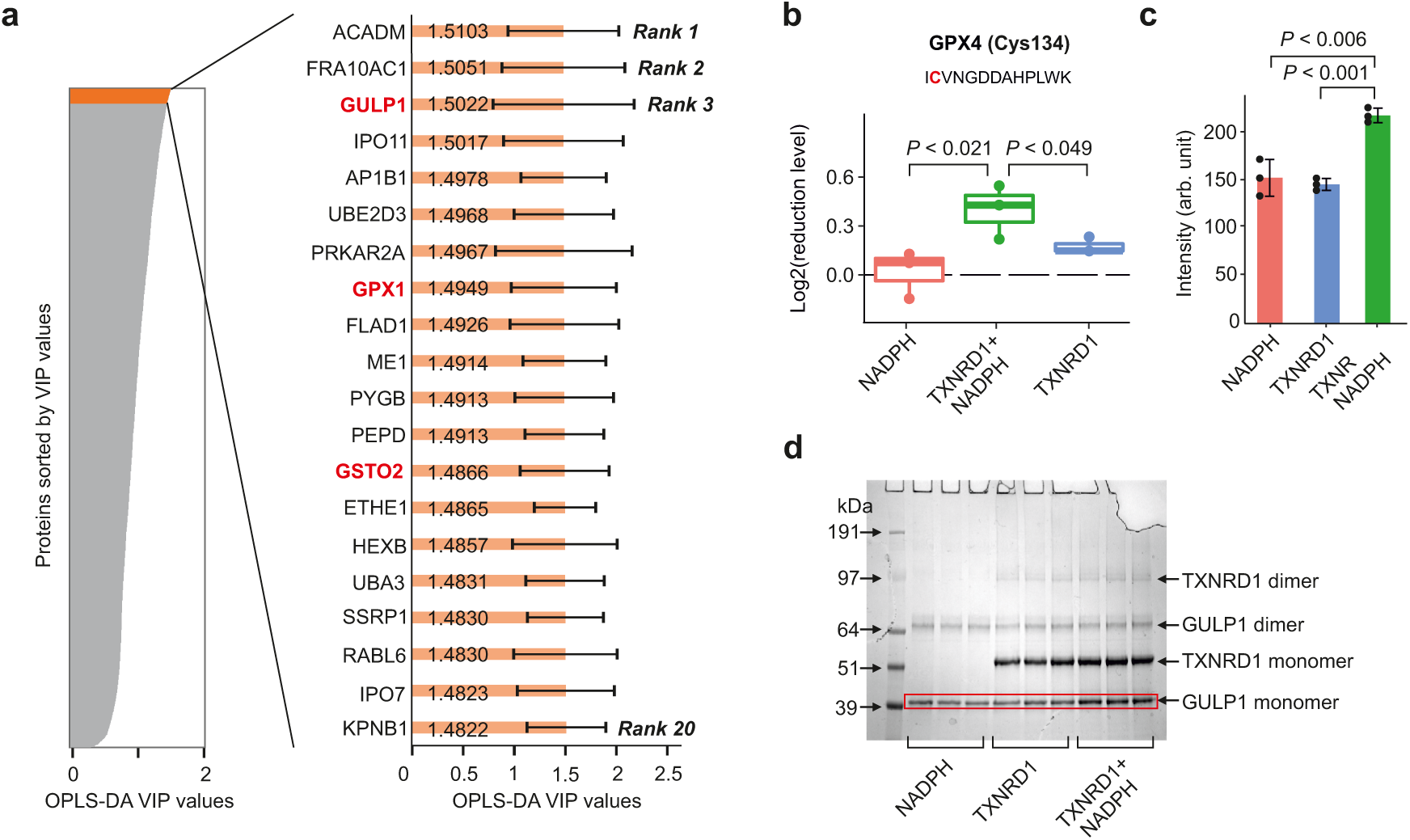
Confirmation of TXNRD1 substrates by orthogonal assays. **a**, The VIP values and confidence intervals (at 95% confidence) extracted from the OPLS-DA model in Fig. 2d. The protein with the highest VIP value is ranked 1^st^ and all the top targets are within the confidence interval. The validated substrates are shown in bold red. **b,** The reduction level of GPX4 Cys134 in the presence of TXNRD1, NADPH or both. **c-d,** The GULP1 monomer levels upon treatment with TXNRD1, NADPH or both (*n*=3 and data presented as mean±SD; two-tailed Student t-test for gel analysis; one-tailed Student t-test for the redox experiment).

**Supplementary Fig. 2.**
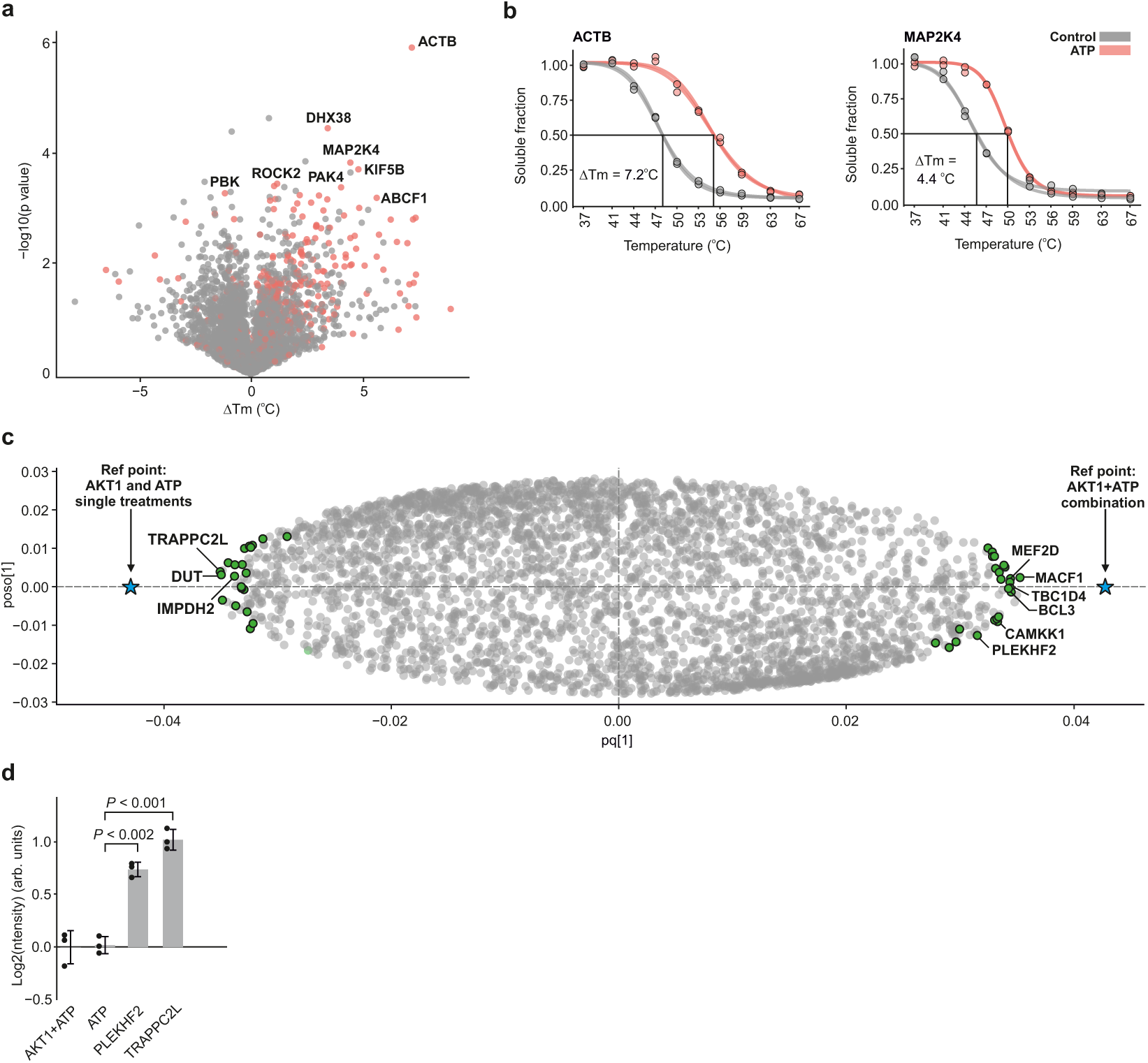
Analysis of SIESTA experiment for AKT1. **a**, Proteins shifting with 500 µM ATP in HELA cell lysate. **b,** Representative proteins shifting with 500 µM ATP. **c,** The OPLS-DA model contrasting the T_m_ in AKT1+ATP treatment against AKT1 and ATP alone. **d,** The relative levels of phosphate release as measured with Phosphoprotein Phosphate Estimation assay (*n*=3 and data presented as mean±SD; two-tailed Student t-test).

**Supplementary Fig. 3.**
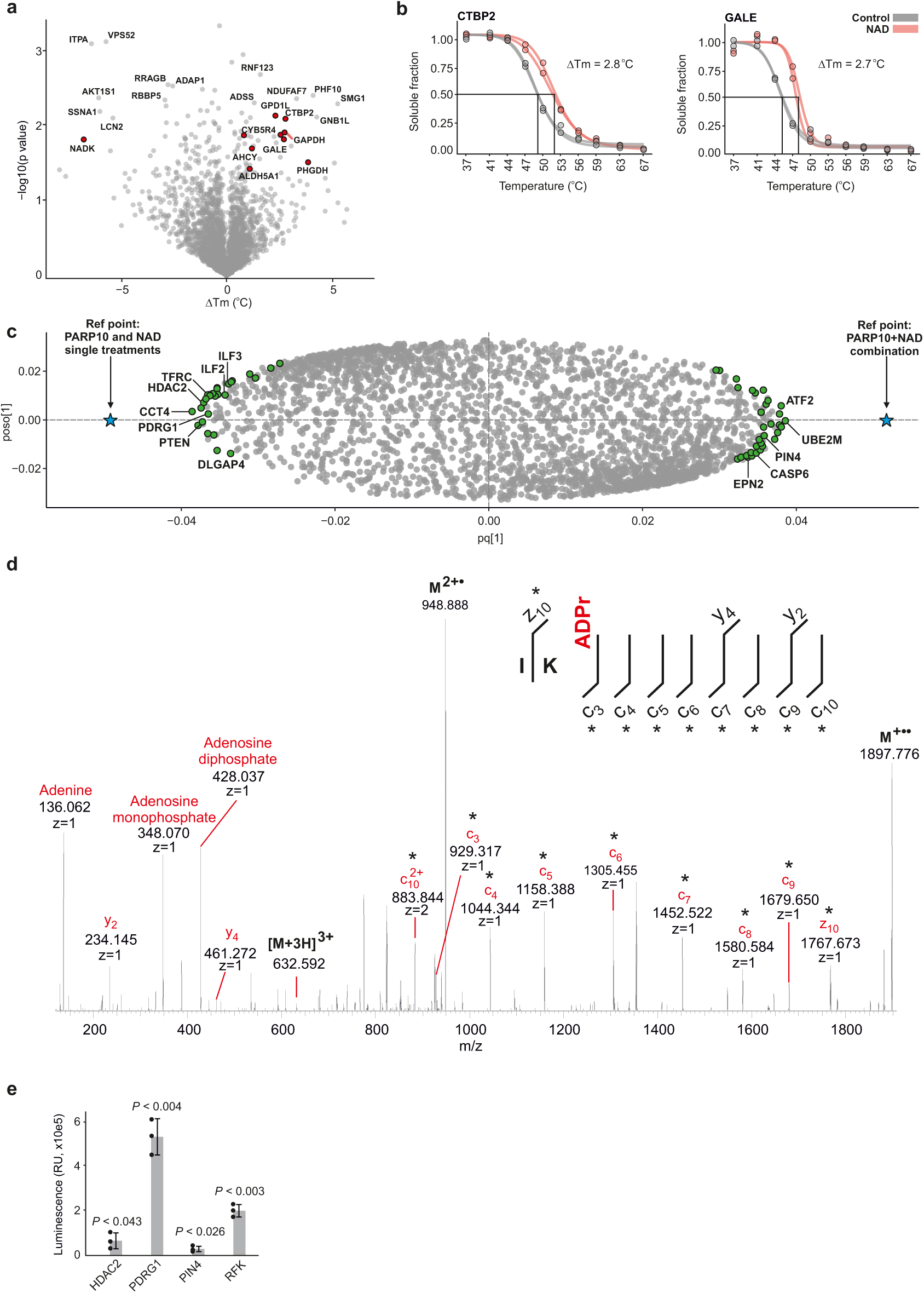
Analysis of SIESTA experiment for PARP10. **a**) A volcano plot with ΔTm of NAD vs. control. The outliers are putative NAD binding proteins (known NAD binders in Uniprot database are shown in red circles). **b,** Representative melting curves of NAD sensor CTBP2 and the NAD binding protein GALE. **c,** Loadings of OPLS-DA model contrasting the “PARP10+NAD” T_m_ vs. those in all other treatments singled out potential substrates (green circles). **d,** Targeted ETD MS/MS of a RFK peptide revealed mono-ADP-ribosylation on glutamic acid residue (the site with the highest sequence-fitting score). The fragments carrying the modification are marked with an asterisk. **e,** The mono-ADP-ribosylation of HDAC2, PIN4, PDRG1 and RFK was confirmed upon incubation with PARP10 catalytic domain and NAD (*n*=3 and data presented as mean±SD; two-tailed Student t-test).

**Supplementary Fig. 4.**
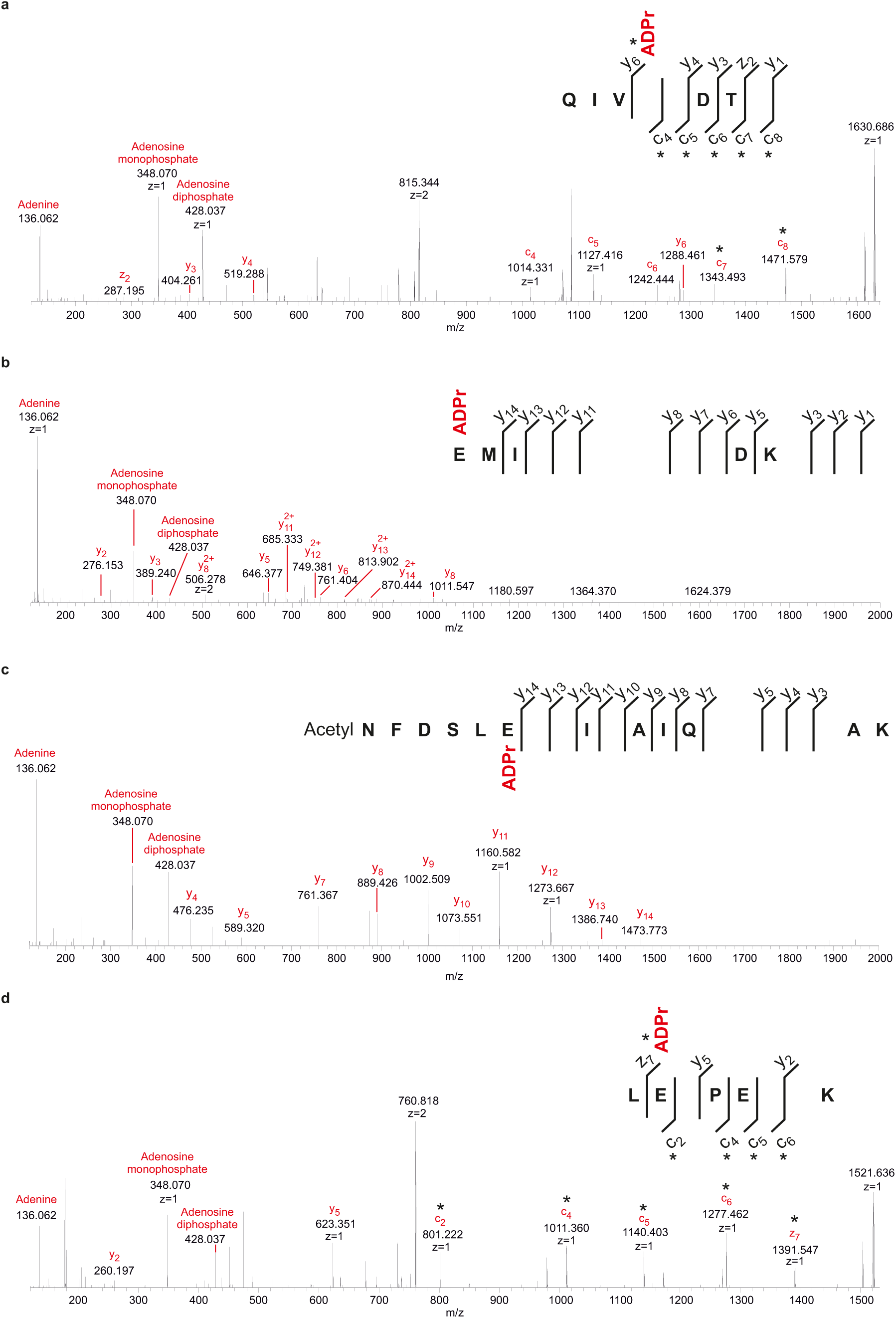
Validation of ADP-ribosylation on multiple sites for PDRG1 and RFK. **a-b**) ADP-ribosylation of PDRG1 and **c-d**) RFK on two sites. Note the presence of signature ions of adenine (*m/z* 136.062), adenosine monophosphate (*m/z* 348.070) and adenosine diphosphate (*m/z* 428.037). The fragments carrying the modification are marked with an asterisk.

**Supplementary Figure 5.**
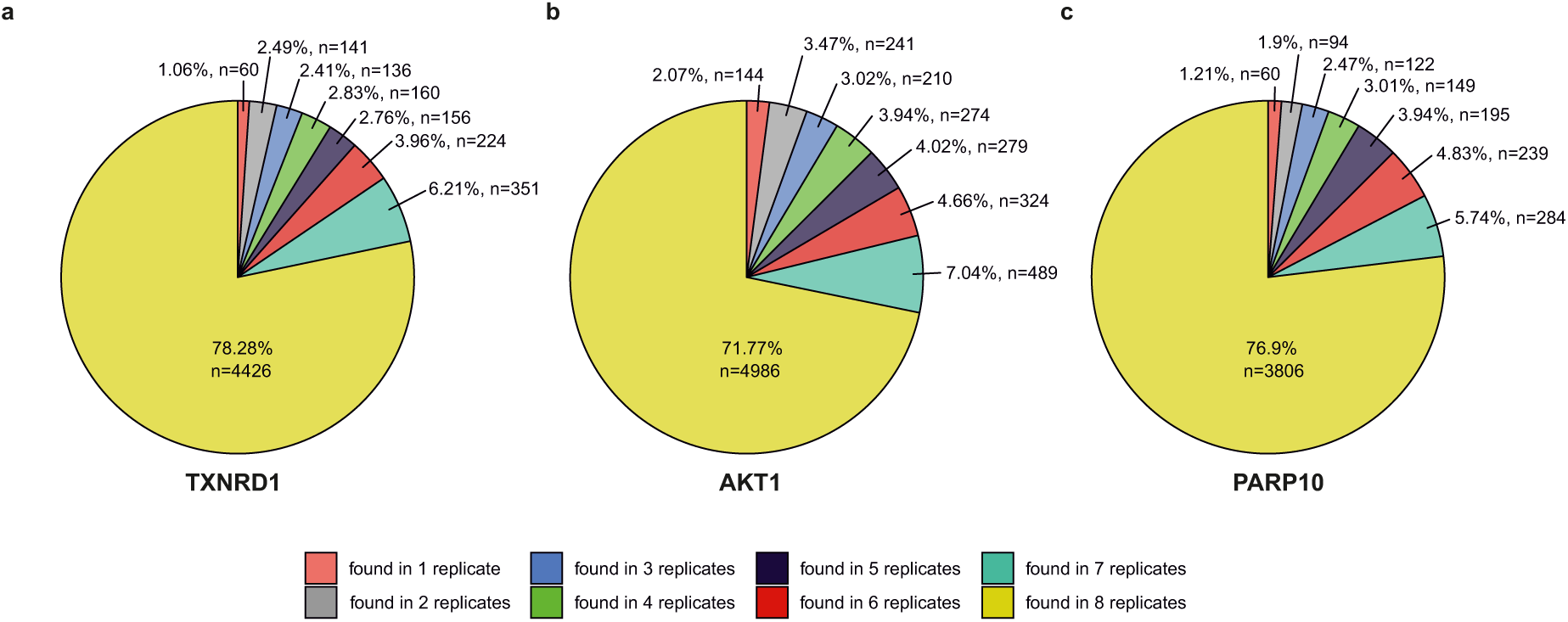
The percentage and number of proteins with at least 2 peptides and no missing values across all replicates in each SIESTA experiment. The useable fraction is shown in mustard.

## Supplementary Tables

**Supplementary Table 1.**
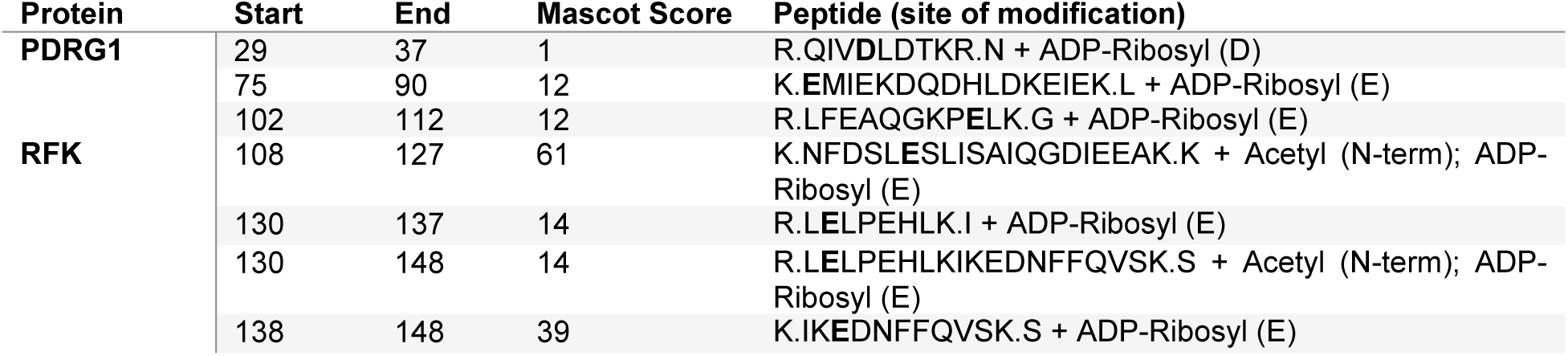
The PARP10-mediated mono-ADP-ribosylation sites of PDRG1 and RFK.

**Supplementary Table 2.** The recombinant proteins used in this study.

| **Protein** | **Catalogue number** | **Source** |
| --- | --- | --- |
| Protein kinase B (AKT1) catalytic domain | 01-401-20N | Carna Biosciences (Japan) |
| RFK | ab89009 | Abcam |
| PDRG1 | PRO-007 | ProSpec |
| Caspase-6 | ALX-201-060-U100 | Enzo |
| HDAC2 | BML-SE533-0050 | Enzo |
| TFRC | 11020H07H50 | Thermo |
| TOX4 | ab211941 | Abcam |
| GSTO1 | NBP1-37093 | Novus |
| GSTO2 | ab124576 | Abcam |
| PRDX2 | ab85331 | Abcam |
| NFKB1 | ab114185 | Abcam |
| GULP1 | ab140546 | Abcam |
| PLEKHF2 | PRO-1874 | ProSpec |
| TRAPPC2L | PRO-1543 | ProSpec |

**Supplementary Table 3.** The LC-MS parameters used in each experiment as well as the number of quantified proteins.

| Ex<br>p.<br>No | Experiment | Cell<br>line | No. of<br>fraction<br>s per<br>replica<br>te | Instrum<br>ent | Total<br>gradient<br>time<br>(min) | MS<br>scan<br>range<br>(m/z) | HCD<br>collision<br>energy | Orbitrap<br>resolution | MS<sup>2</sup><br>resolution | MS<br>AGC<br>target | MS<sup>2</sup><br>AGC<br>target | MS<br>max<br>injection<br>time<br>(ms) | MS<sup>2</sup><br>max<br>injection<br>time | Isolation<br>window | Dynamical<br>exclusion | No. of<br>proteins | No. of<br>proteins<br>with at<br>least 2<br>peptides |
| --- | --- | --- | --- | --- | --- | --- | --- | --- | --- | --- | --- | --- | --- | --- | --- | --- | --- |
| 1 | SIESTA<br>TXNRD1 | HCT1<br>16 | 8 | Fusion | 120 | 400-160<br>0 | 40 | 120,000 | 60,000 | 1e6 | 1e5 | 50 | 105 | 0.7 | 60 | 5864 | 5637 |
| 2 | SIESTA<br>PARP10 | HCT1<br>16 | 8 | Fusion | 160 | 400-160<br>0 | 40 | 120,000 | 60,000 | 1e6 | 1e5 | 50 | 105 | 0.7 | 60 | 5194 | 4979 |
| 3 | SIESTA<br>AKT1 | HELA | 24 | Q<br>Exactive<br>HF | 95 | 375-150<br>0 | 33 | 120,000 | 45,000 | 3e6 | 2e5 | 100 | 120 | 1.6 | 45 | 7179 | 6918 |
| 4 | Redox<br>experiment<br>with<br>iodoTMT | - | - | Q<br>Exactive | 120 | 375-150<br>0 | 32 | 70,000 | 35,000 | 3e6 | 2e5 | 120 | 120 | 1.2 | 45 | - | - |

